# Inverted Face with Upright Body: Evidence for Face Inversion Effect in Japanese Macaques (*Macaca fuscata*) under the Preferential Looking Procedure

**DOI:** 10.1101/266676

**Authors:** Masaki Tomonaga

## Abstract

Young Japanese macaques (*Macaca fuscata*) looked at the photographs that contained faces of the two species of macaques (*M. fuscata* and *M. mulatta*) for longer time when they were presented at upright orientation than inverted orientation, and that species discrimination was deteriorated when the photographs were presented at inverted orientation [16]. The present experiment further explored the factor affecting the inversion effect observed in the previous experiments. Five young laboratory-born Japanese macaques looked at the photographs by pressing the lever under the conjugate schedule of sensory reinforcement, that is, successive preferential looking procedure. Photographs of macaques were scrambled and made up two types of the bizarre photographs: upright face with upright body (orientation consistent) and inverted face with upright body (orientation inconsistent). These two types of photographs were presented at upright and inverted orientations. The monkeys looked longer when the face was upright than inverted. Orientation of body and background had no effect on looking time duration. The present results strongly suggest that inversion effect found in macaques is face-specific.

## Introduction

Face perception by humans is considered different from other visual perception in that there is severe impairment of perception (identification and discrimination) when faces are presented with inverted orientation. This phenomenon is called “inversion effect” [1, 19]. On the contrary, the face inversion effect in other nonhuman animals, especially primates, was still controversial. Nonhuman primates showed evidence for inversion effect as well as against it. Advances in this topic, however, accumulated evidence *for* face inversion effect. When using the large number of photographs, nonhuman primates show the face-specific inversion effect [13, 14, 16, 17, 18]. For example, Tomonaga [17] tested the inversion effect in a chimpanzee using the identity and rotational matching tasks with 104 human faces. He found the face-specific inversion effect in the chimpanzee when using a large number of examples. Parr et al. [13] also found similar finding in chimpanzees. Tomonaga [16] examined face inversion effect in laboratory-raised Japanese macaques (*Macaca fuscata*) under the modified preferential looking task. Monkeys were allowed to look at the photographs during they held down the lever. Monkeys showed significant difference in preference assessed with looking time between upright rhesus and Japanese macaque photographs [4], whereas no difference when photographs were presented at 90 or 180 degrees. This inversion effect was the strongest for the photographs containing front faces clearly. When photographs contained no clear faces, the inversion effect disappeared.

In his experiment, however, orientations of faces, bodies, and backgrounds were changed together, so that, it is still unclear what was the source of inversion effect. It might be possible that the monkeys utilized other cues than facial orientation for recognition of photographs. In the present experiment, the monkeys were presented a set of scrambled photographs of macaques. Each photograph contained a single monkey with the same orientation of face, body, and background, or inconsistent orientation between face and body. By using these scrambled photographs, it can be directly tested which orientation is critical for inversion effect in monkeys, faces or other parts of photographs. If the face orientation were critical, they would show inversion effect based on face orientation irrespective of the types of photographs.

## Method

### Participants

The participants were the five young Japanese macaques (*Macaca fuscata*). At the onset of the present experiments, they were 5 years old in average (range: 4 to 6). They had been isolated from their mothers within one week after birth and raised by human caretakers. They lived in a cage (70 × 70 × 70 cm) with another macaque. They have had an extensive history in participating the experiments using the modified preferential looking procedure as in the present experiments [4, 16]. During the present study, they were not deprived of either food or water. This study was approved by the Animal Welfare and Care Committee of the Primate Research Institute, Kyoto University, and followed Guidelines for the Care and Use of Laboratory Primates of the Primate Research Institute, Kyoto University

### Apparatus

The apparatus was the same as the previous studies [4, 16]. Two identical chambers (60 × 60 × 60 cm) were set up in the experimental room. The front panel (40 × 40 cm)of each chamber was made of clear Plexiglass. A slide projector (CABIN, model Family Cabin) was installed behind the screen and projected the slide photographs onto a 33 × 33 cm opaque screen placed 50 cm apart from the front panel. A single lever with a red lamp was installed in the lower center of the front panel. All experimental events were controlled and recorded by MSX2 personal computers (TOSHIBA, model HX-34).

### Stimuli

Four types of color photographs of macaques (Japanese and rhesus macaques) were prepared. Each photograph contained a single individual and a full face taken in front. Figure 1 shows examples of four types of photographs. Each photograph was based on an original photograph. In the orientation consistent groups, the part of face was cut and pasted at a slightly shifted position. In the orientation inconsistent groups, the part of face was cut, inverted, and then pasted. These photographs were presented at the two orientations, upright and inverted, so yielded four types of photographs. Twenty original photographs were prepared, so that, each monkey was shown 160 pictures, and additional control slides; 20 no-light slides and 20 white-light slides. These 200 slides were set in the two circular slide cartridges, which had 100 slots.

**Figure 1.**
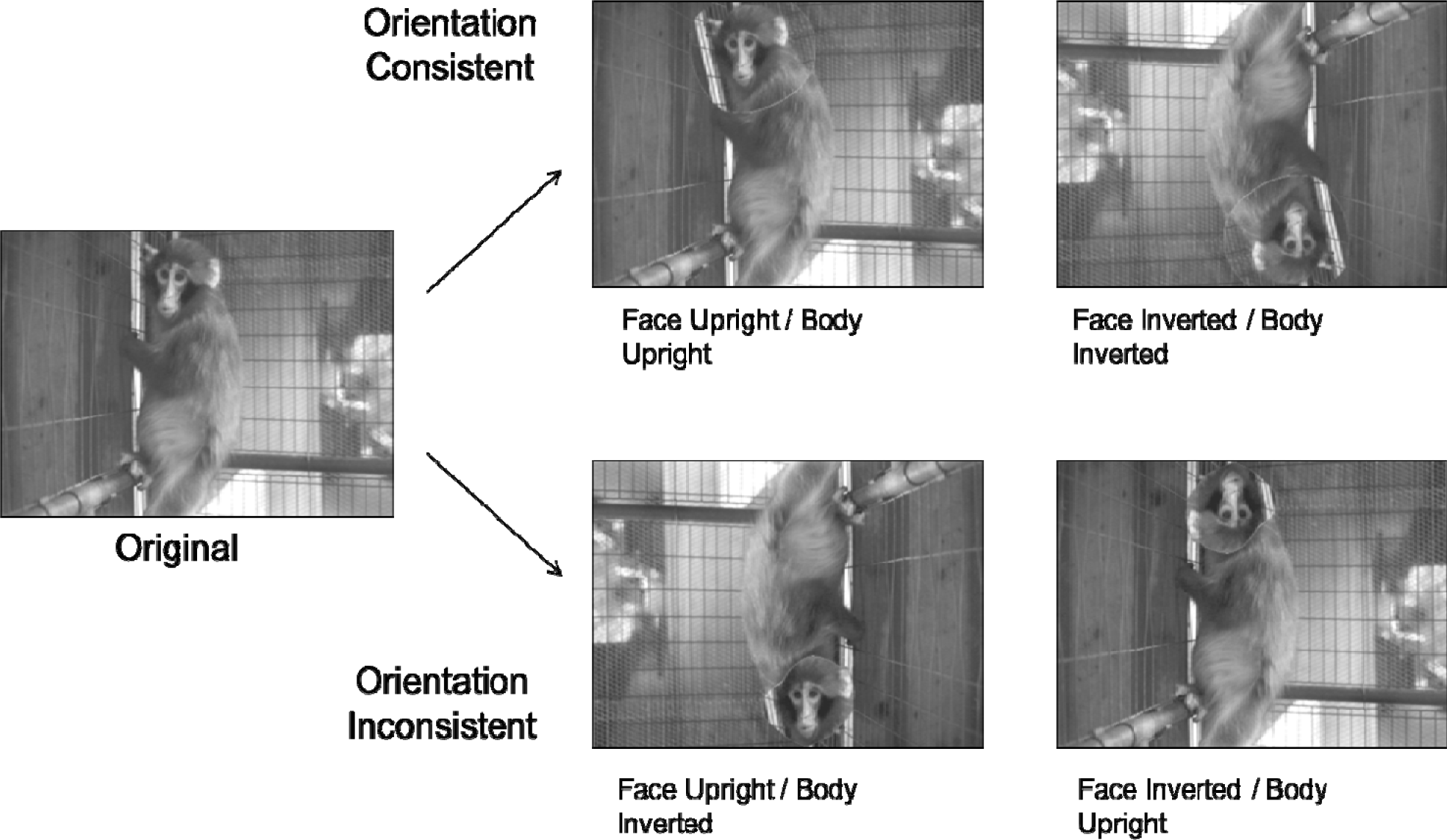
Examples of four types of photographs used in the present experiment.

### Procedure

Figure 2 represents the schematic diagram of the procedure. In the present experiments, the modified version of preferential looking task was employed [3, 4, 5, 16]. When the monkey held down the lever in the presence of red light, the slide was projected onto the screen during the monkey’s holding of the lever. If the monkey released the lever or the lever holding exceeded 10 s, the slide was terminated. Very short response duration (faster than 100 ms) had no effect. When the monkey held down the lever again within the 10 s from the last lever release, the same slide was presented again. On the other hand, if the monkey did not press the lever more than 10 s, the next slide was set up for presentation. This interval was called interresponse interval. The trial was defined as the presentations of the same slide. Since the slide presentation was contingent upon the monkey’s response, it was procedurally considered as sensory reinforcement [5]. Since the duration of the trial was dependent upon the monkey’s responses, this task was also considered as “successive” preferential looking task. The strength of sensory reinforcement, or preference index for each slide was identified by measuring the response duration (D) and interresponse interval (I). The index was calculated with the formula below.

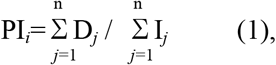

where PI_*i*_, designated the preference index of the *i*-th trial, n designated the number of responses in the *i*-th trial, D*_j_* designated the response duration of the *j*-th response, and I*_j_* designated the interresponse interval after the *j*-th response. When this index was higher, the monkey looked at the photograph for longer time and looked again with shorter interval. Because of the between-paricipants variances of the preference indices, the normalized index for each monkey calculated with the formula (2) was used for the data analyses.

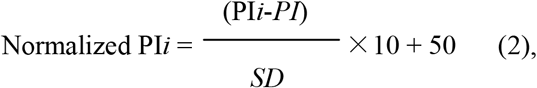

where the Pl*i* designated the preference index for the slide *i*, *PI* and *SD* designated the mean and standard deviation of the preference indices for all slides. By normalizing, the mean value of preference index for each monkey was equated to 50.

**Figure 2.**
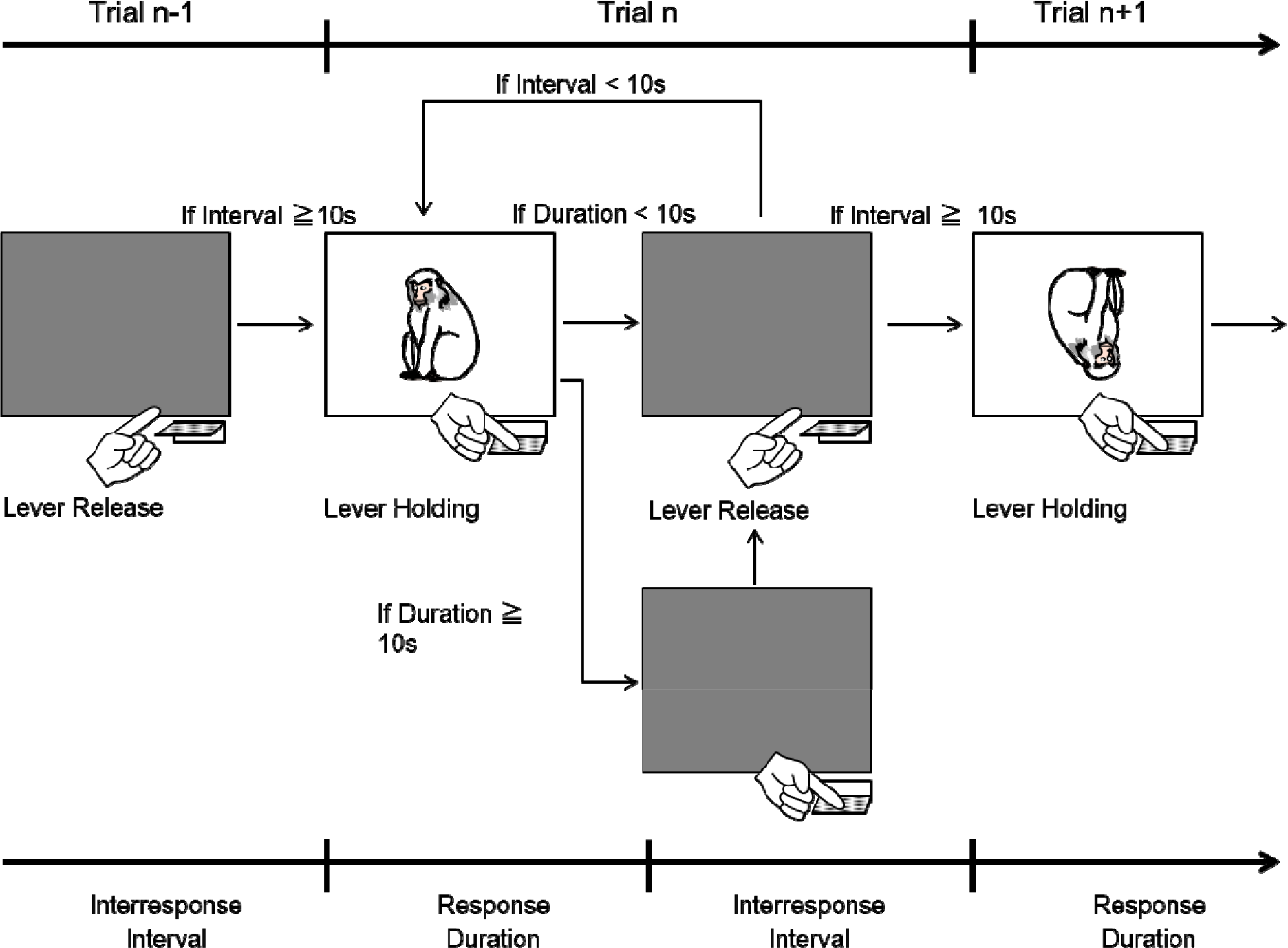
Schematic diagram of the successive looking time experiment.

Each slide was presented five times. The preference index for each slide was averaged across these five trials. These indices were normalized and then averaged for each type of photographs. A session was continued until the monkey completed 500 trials, that is five rotations of one cartridge. Each monkey received for two sessions (i.e. one session for one cartridge). The duration of each session was approximately 6 hours.

## Results

Figure 3 shows the normalized preference index for each type of the photographs averaged across monkeys. The preference indices of control slides were 39.98 for no-light slides and 44.42 for white-light slides. These values were higher than those for photograph slides (51.49 in total). As shown in the figure, the monkeys showed higher preference index when the photographs contained upright face than when it contained inverted face, irrespective of body and background orientations. Normalized preference indices were used for a repeated-measures two-way analysis of variance (ANOVA), in which the Face orientation (upright vs. inverted) and Type of photographs (orientation consistent vs. inconsistent) were within-participant factors. The results of ANOVA showed the significant main effect of Face orientation [*F*(1, 3)=21.85, *p*<0.01], but no effect of Type of photographs [*F*(1, 4)=4.92, *p*=0.091] and interaction [*F*(1, 4)=0.20].

**Figure 3.**
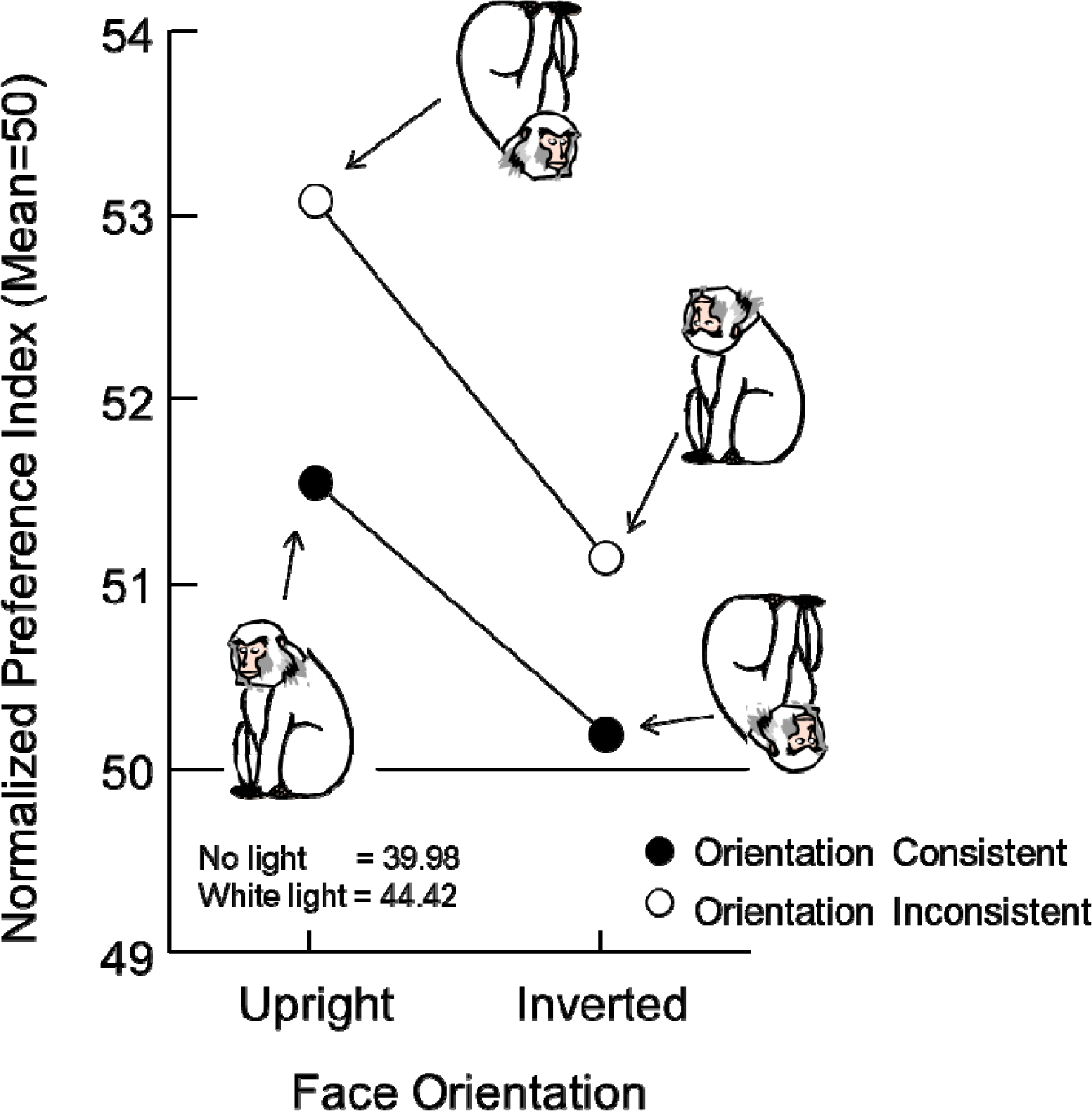
Mean normalized preference index for each type of photographs.

## Discussion

The results of orientation-consistent photographs well replicated those of the previous experiments using the same monkeys and the same procedure [16]. The monkeys showed higher value of the preference index when the face was presented at upright orientation. When the orientation-inconsistent photographs were presented, they also looked at the photographs with upright faces. These results clearly indicate that the inversion effect was mainly controlled by the orientation of faces but not by the body and background orientations. The inversion effect is face-specific.

In the study with macaques, they showed inversion effect only for human faces but not for monkey faces and scenery photographs [18]. These results were discussed on the basis of difference in discriminability between human and monkey faces. The extraneous or uncontrolled factors in the photographs of monkey faces such as homogeneity, species, face angle, background, and so on, might have affected on the inversion effect. The present and previous experiments [16], however, clearly showed the face-inversion effect for monkey faces. In the present experiment, the variability among photographs can not explain the present results. The photographs used in the present experiment were as variable as those used in the Wright and Roberts’ study [18] in respect to face angle and background. Wright and Roberts used photographs of various primate species. As [13] suggested, species-level or genus-level homogeneity might cause inversion effect in the present experiment.

Upright faces and inverted faces are processed in different ways in humans. Upright faces are processed on the basis of configural properties of faces, while the inverted faces are processed on the basis of local features [2]. It might be possible that the difficulty in recognizing inverted faces might be due to the degradation of configural information in monkeys as well as in humans.

In humans, face perception and face-inversion effect are specialized in right hemisphere [7, 8, 9, 10, 15, 20]. For monkeys, Hamilton and Vermeire [6] reported the right hemispheric advantage in face perception in split-brain macaques on the one hand, while Overman and Doty [12] failed to demonstrate hemispheric specialization in macaques, on the other hand. In chimpanzees, right hemispheric advantage were reported by using chimeric face presentations [11, 17]. We should further examine the relationship between face perception and functional cerebral asymmetry in nonhuman primates.

In summary, the present results verified the inversion effect in face perception in macaques under the preferential-looking task. This effect was controlled only by orientation of faces but not by other background orientations. Inversion effect is face-specific also in macaques.

## Acknowledgments

This study was financially supported by the JSPS/MEXT KAKENHI (#4710053, #13610086, #20002001, #23220006, #24000001, #15H05709, #16H06283), JSPS-CCSN and the JSPS Leading Graduate Program in Primatology and Wildlife Science (U04) at Kyoto University. The author wishes to thank Drs. Kazuo Fujita and Tetsuro Matsuzawa, and the staff of the Primate Research Institute, Kyoto University for their invaluable comments on this study.

